# A framework for assessing the skill and value of operational recruitment forecasts

**DOI:** 10.1101/2021.07.05.451182

**Authors:** Christian Kiaer, Stefan Neuenfeldt, Mark R. Payne

## Abstract

Forecasting variation in the recruitment to fish stocks is one of the most challenging and long-running problems in fisheries science and essentially remains unsolved today. Traditionally recruitment forecasts are developed and evaluated based on explanatory and goodness-of-fit approaches that do not reflect their ability to predict beyond the data on which they were developed. Here we propose a new generic framework that allows the skill and value of recruitment forecasts to be assessed in a manner that is relevant to their potential use in an operational setting. We assess forecast skill based on predictive power using a retrospective forecasting approach inspired by meterology, and emphasise the importance of assessing these forecasts relative to a baseline. We quantify the value of these forecasts using an economic cost-loss decision model that is directly relevant to many forecast users. We demonstrate this framework using four stocks of lesser sandeel (*Ammodytes marinus*) in the North Sea, showing for the first time in an operationally realistic setting that skilful and valuable forecasts are feasible in two of these areas. This result shows the ability to produce valuable short-term recruitment forecasts, and highlights the need to revisit our approach to and understanding of recruitment forecasting.

## Introduction

Recent developments in ocean observations and modelling today make it possible to forecast many of the physical variables in the ocean (Doblas-Reyes *et al.*, 2013; Meehl *et al.*, 2014). Building on top of this data about the ocean environment, forecasts of marine ecological responses have been developed (Payne et al 2017) and provide managers and stakeholders the foresight needed to sustainably manage marine living resources (Tommasi *et al.*, 2017b; Hobday *et al.*, 2018). Examples of operational forecasts already in use include southern Bluefin tuna habitat forecasts (Eveson *et al.*, 2015), the dynamic fisheries bycatch management tool EcoCast (Hazen *et al.*, 2018), and blue whale habitat preference forecast (Hazen *et al.*, 2017). However, these operational fisheries forecast products are currently limited to predictions of distribution and phenology and there are currently no known operational marine fish recruitment forecasts (Payne *et al.*, 2017).

Understanding and forecasting changes in fish stock productivity has, however, been a key aspiration in fisheries science for the last century (Leggett and Deblois, 1994; Subbey *et al.*, 2014; Tommasi *et al.*, 2017a; Haltuch *et al.*, 2019). Recruitment, the number of young individuals produced each year, has a key role in shaping fish population dynamics (Hilborn and Walters, 1992), especially in determining total allowable catches for short-lived species, where the recruiting year-classes contribute a significant share of the landings. Environmental drivers play an important role in shaping the productivity of such stocks (e.g. via temperature (MacKenzie *et al.*, 2008; Mantzouni and Mackenzie, 2010), salinity (Köster *et al.*, 2005) or phenology (Platt *et al.*, 2003)) and including climate information in stock-assessments can reduce uncertainties in stock status and the risk of over- or under harvesting (Hare *et al.*, 2010; Haltuch and Punt, 2011; Tommasi *et al.*, 2017a, 2017b). The ability to foresee changes in productivity on a short time-scale can therefore enable adaptive and pre-emptive decision-making strategies, benefiting both stakeholders and managers (Hobday et al., 2016; Payne et al., 2017; Welch et al., 2019).

Common approaches have however shown limited ability to produce reliable recruitment forecasts for operational (i.e. regularly repeated) use in management. The large variety of underlying environmental, physical and ecosystem processes affecting recruitment simultaneously (Leggett and Deblois, 1994; Browman *et al.*, 1995; Myers, 1998; Tommasi *et al.*, 2017b) can often give rise to transient but spurious correlations (Sugihara *et al.*, 2012). Fish population time series are often relatively short in length (Ricard *et al.*, 2012) and hampered by high observation noise, limiting the ability to develop and test predictive models (Clark and Bjørnstad, 2004; Ward *et al.*, 2014). Furthermore, environment-recruitment correlations have been shown to breakdown when confronted with new data, diminishing the uses for management (Myers, 1998; Tommasi *et al.*, 2017b). The relative importance of drivers of recruitment can also change from year to year (“non-stationarity”) (Subbey *et al.*, 2014; Haltuch *et al.*, 2019). As a consequence of all of these processes, recruitment forecasts are widely viewed with scepticism in the community today.

Nevertheless, the potential of such forecasts to benefit all those that depend on living marine resources is clear. So how can this potential be realised? And even more importantly, how would we know when we have produced forecasts that can be used as a regular part of decision-making? To answer this question, here we take inspiration from other forecasting fields, and in particular from meteorology, a discipline that has also been attempting to predict chaotic and difficult to observe systems for nearly a century (albeit with considerably more success!). In particular, the question of “what makes a good forecast?” is addressed in a seminal 1993 paper in the field by Alan Murphy (Murphy, 1993) that introduces two key relevant concepts, skill, and value, which form the basis for this work.

Murphy defines forecast “skill” as the quantitative ability of the forecast: is it numerically correct? In the marine setting, model performance is often measured based on goodness-of-fit measures that quantify the ability to explain the data e.g (Lindegren *et al.*, 2018). There is however, a fundamental difference between explanatory and predictive power: while *explanatory* models can be used to investigate causal hypotheses, models with high explanatory power cannot be expected to predict well (Levins, 1966; Shmueli, 2009; Dickey-Collas et al., 2014). But when the goal is to produce forecasts to be used regularly to predict into the future for use in a decision-making context, we clearly need to evaluate their *predictive* power. In the atmospheric and climate sciences for example, skill is often assessed based on retrospective forecast analysis (Wilks, 2011) i.e. predicting beyond the period over which the model was developed or tuned, directly reflecting the way the forecast would be used operationally. Furthermore, meteorology always places its forecasts in the context of a baseline or reference forecast (Jolliffe and Stephenson, 2012; Payne *et al.*, 2012). Common baseline forecasts includes random selection of categories or using the average over a given reference time period, often referred to as climatology in atmospheric sciences (Jolliffe and Stephenson, 2012).

Secondly, Murphy discusses the usefulness of a forecast in terms of its “value” in aiding decision-making. A good forecast is of value to an end-user by assisting in decision-making, providing economic value or otherwise benefiting the user (Murphy, 1993). While value in recruitment forecasts has been discussed (e.g. Walters, 1989; Field *et al.*, 2010), a quantitative approach to value is rarely seen in marine science. Simple economic decision models can analyse forecasts under simplified assumptions, helping end-users decide if it is economically wise to follow the forecast (Murphy, 1976a). Quantitatively providing a value assessment can help integrate forecast products directly into a user’s framework, allowing users to assess the benefits of a given forecast system and can give a clear insight into how, and when, a forecast should be used (Murphy, 1976a).

Here we argue that as the recruitment problem has never been evaluated from this perspective before, we currently do not know whether it is possible to regularly make skilful and valuable forecasts of recruitment. We therefore combine the ideas Murphy (1993) with the state of the art in recruitment modelling to give a generic framework for developing and assessing short-term recruitment forecasts for fish stocks for regular use in an decision-making setting. Forecast skill is assessed based on predictive performance, using validation techniques currently used in atmospheric and meteorological sciences and that reflect the way a forecast would be used in practice. Value is assessed quantitively, using an economic cost-loss decision model, providing insight into the actual monetary value of the forecast product. We demonstrate the framework using multiple stocks of the ecologically and economically important lesser sandeel (*Ammodytes marinus*) in the North Sea, where previous studies of recruitment have already highlighted several recruitment correlates (Arnott and Ruxton, 2002; van Deurs *et al.*, 2009; Lindegren *et al.*, 2018).

## Methods

### Recruitment forecast framework

This work presents a generic framework (*F*igure *1*a) for assessing recruitment forecasts of fish stocks in an operational setting. The core of the framework is the idea of retrospective forecasting, an approach adapted from the atmospheric sciences, in which the time series of interest is split into two continuous blocks either side of a hypothetical “forecast issue date”. The first block is used to parameterise and train the core predictive model (the “training” block): predictions are then made for the remaining block of data (the “verification” block) based on this model. The issue date is then shifted forward by one time step, the data repartitioned and the process repeated. Iterating over all issue dates, a database of predictions is generated, with each prediction being characterised by the id of the cohort being predicted and issue date: the difference between these two is the “lead time” of the forecast (*F*igure *1*b). The ensemble of predictions can then be compared against the “true” recruitment to that cohort, with various skill metrics being calculated as a function of forecast lead time. The skill metrics generated are then used as the basis for forecast value assessment.

**Figure 1.**
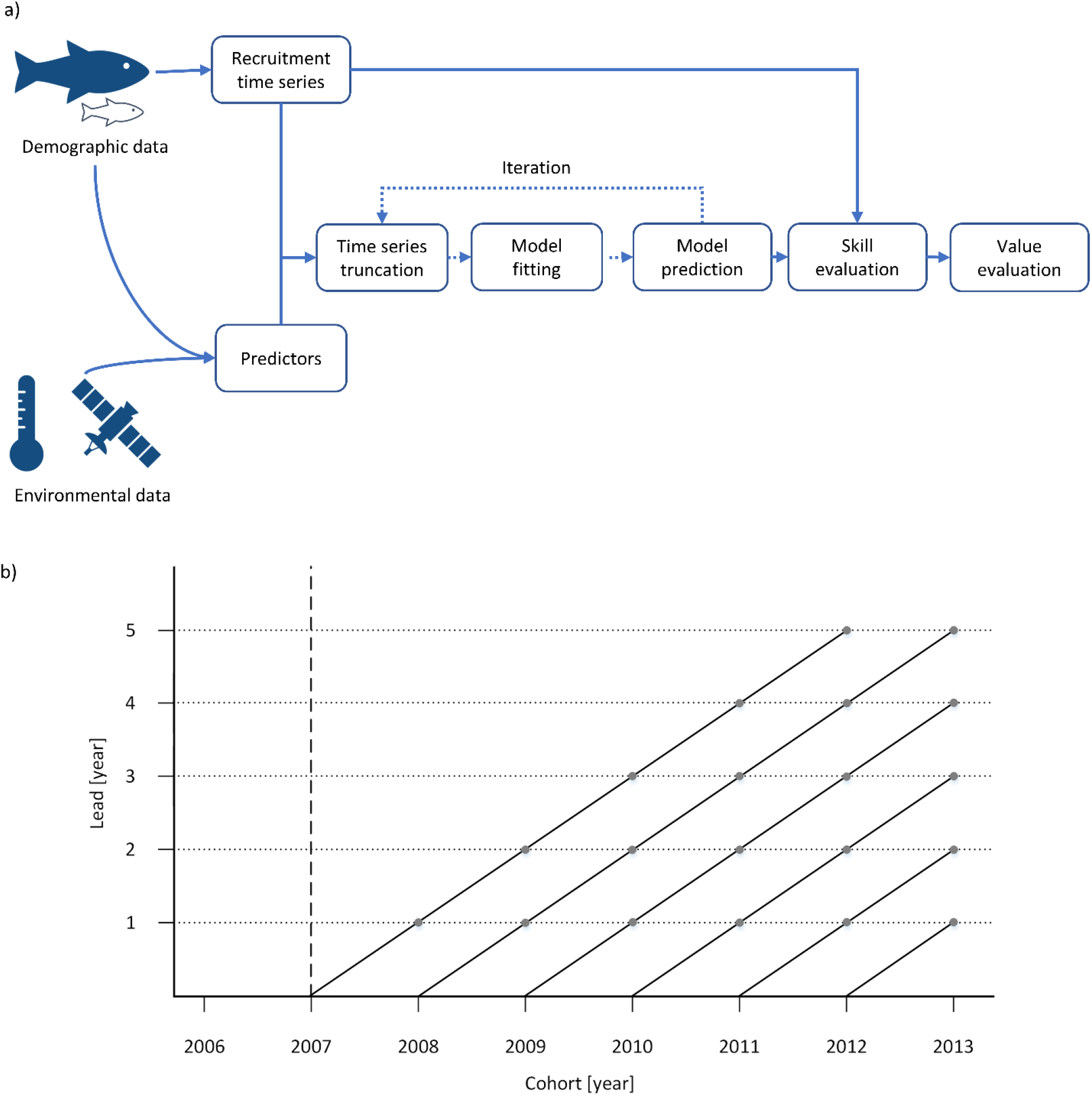
Skill and value assessment framework. a) Overview of the process in the forecasting framework. Here, data are extracted and combined into the appropriate modelling data. Afterwards, an iterative process of data truncation and model fitting are the basis of all model objects and predictions. These predictions contains both the current and the retrospective predictions, which can be used for skill and value evaluation b) Schematic of retrospective forecast system used to generate a retrospective forecast time series. One time series is generated at each lead time. Dashed line indicates first data cut-off and the start of the retrospective forecasting period. Dotted lines indicates the forecast time series at a given lead. After the first cut-off each subsequent retrospective forecast will include the previous year’s observations increasing the size of the model training data set. For each generated retrospective forecast time series skill, value and accuracy will be evaluated. Depending on species, stocks, data availability and period of interest, the evaluated cohort period and the start of the retrospective analysis can vary.

There are several key features of this framework that make it highly appropriate for addressing the question at hand i.e. assessing operational forecast skill. The emphasis on temporal blocks, for example, differs from other cross-validation approaches (of which it is a subset (Roberts *et al.*, 2017)) and is important as it directly mimics the way in which recruitment forecasts would be used in an operational setting. Furthermore more, temporal blocks also remove the potential for the leakage of information between randomly-selected cross-validation folds, a particularly important issue where there is temporal structure and autocorrelation in the the time series (as is common in recruitment data). This retrospective forecasting approach therefore gives a much more realistic assessment of the skill of forecast, and has been shown to consistently outperform other approaches when forecasting is the goal (Roberts *et al.*, 2017).

The user of the temporal-block approach, however, has two key caveats associated with it. Firstly, the choice of the initial forecast issue date separating the training and verification blocks represents a tradeoff between the desire to have as many verifications as possible (and thus the most reliable skill evaluation) and the need to have sufficient data to train the model on in the first place. This tradeoff is more restrictive than random cross-validation and will be particularly acute in instances where the length of the time-series is short: in some cases, there may not be sufficient data to make a reliable skill assessment in this manner. The exact choice will depend on the characteristics of the system at hand. Secondly, and even more importantly, care must be taken to avoid inadvertedly introducing circular reasoning through the use of predictors identified by explanatory analyses over the whole time series: such variables will show skill over the length of the time-series for which they were indentified, but this may not extend into the future. Ideally, predictors should be based on either generic reasoning (e.g. stock-recruitment relationships, the match-mismatch hypothesis) or work published prior to the earliest forecast issue date considered. Alternatively, automatic variable and/or model selection procedures can be incorporated into the “fit model” part of the framework to allow the identification of skilful predictors for each forecast issue date.

The generic nature of the framework mans it can be applied widely: each individual application can and should vary depending on the specifics of the system being assessed. The recruitment time series used can be taken from either stock assessment outputs or from a recruitment-index (e.g. from a larval survey). The selection of predictors is flexible but should be informed by the best available biological knowledge about the stock (Dickey-Collas *et al.*, 2014b; Subbey *et al.*, 2014) (previous caveats not withstanding): stock-specific biomass or demographic indicators, environmental data or other biological parameters (e.g. prey and predator concentrations) can be incorporated equally. Any modelling approach that produces predictions can be considered, including classical recruitment models (e.g. Ricker (Ricker, 1954) and Beverton-Holt (Beverton and Holt, 1957)), statistical and data mining approaches (e.g. generalized additive models (GAMs) (Hastie and Tibshirani, 1986), empirical dynamic modelling (EDM) (Sugihara et al., 2012) and classifier models (Fernandes et al., 2015)): ensembles of models can also be considered e.g. combined via multi-model inference (Burnham and Anderson, 2004). Predictions can (and should) be considered in terms of continuous outputs, probability distributions and/or as categories (i.e.. using a division into terciles (high, medium, low) based on historical observations). The choice of skill metrics will be influenced by the nature of the forecast (Jolliffe and Stephenson, 2012) but should include multiple metrics(Stow *et al.*, 2009; Brun *et al.*, 2016). Skill metrics then form the basis for a quantitative value assessment, evaluating the expected economic value of following a given forecast. Furthermore, the framework allows for forecasts of both single stocks or of aggregations of multiple stocks into a single portfolio forecast, as may be relevant for decision-making across wider-scales (e.g. factories processing many different species)

We illustrate the use of this framework through a worked example focusing on recruitment forecasts of the lesser sandeel (*Ammodytes marinus*) in the North Sea below.

### Sandeel Case Study

The lesser sandeel is a pelagic species of the Ammodytidea family and is one of the most common sandeels found in the North Sea. Adult lesser sandeel habitats are found in most of the North Sea, generally distributed across shallow sandy banks (van Deurs et al., 2009, Figure 1a). Particle-tracking studies and the sedentary state of post-recruitment sandeel (Christensen et al., 2008; Pedersen et al., 2019) resulted in a division into 7 different individually managed North Sea sandeel stocks. Analytical stock assessments are done in management areas 1r, 2r, 3r and 4 (see Figure 2a), while the remaining three stocks are considered data poor. Sandeel is seen as one of the main links between primary production and the higher trophic levels in the North Sea for both larger piscivorous fish (e.g. cod and haddock) and seabirds (Eliasen *et al.*, 2011). The lesser sandeel has historically supported a large fishery, which has seen a large decline in recent years (Dickey-Collas *et al.*, 2014b). Due to the importance of the species, recruitment to these stocks is well studied (Arnott and Ruxton, 2002; van Deurs *et al.*, 2009; Eigaard *et al.*, 2014; Lindegren *et al.*, 2018). In the south western part of the North Sea (i.e. management area 1r), sandeel shows signs of being influenced negatively by temperature, while the abundance of the main prey, *Calanus finmarchus*, has a positive influence (Arnott and Ruxton, 2002; Lindegren *et al.*, 2018). Density dependence has also been found to be an important driver, where competition with young adults and juveniles has a negative effect on recruitment (van Deurs *et al.*, 2009). Currently, stock assessment uses a geometric mean for recruitment predictions (ICES, 2018). These geometric means will be used as continuous reference models during skill evaluation.

**Figure 2.**
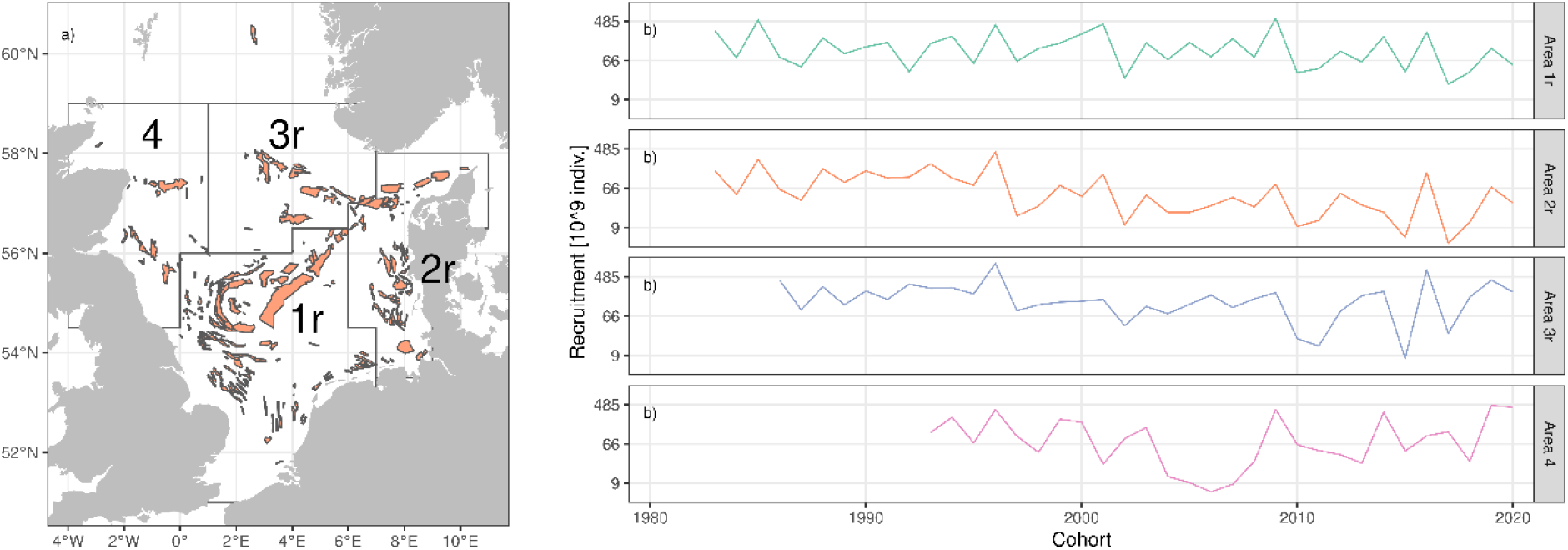
Study area and data. a) Map of the North Sea showing the four management areas of sandeel assessed analytically. Sandy habitat banks, the predominant sandeel habitat, are shown in orange. b) Recruitment time series for the four sandeel stocks from the official ICES stock assessment. Dashed-horizontal lines mark the delineation of the upper and lower terciles for each stock.

### Data

Operational forecasts require data to be available at the time of the forecast, potentially excluding some potentially relevant predictors. For example, estimates of zooplankton prey, *Calanus finmarchicus* and *Temora longicornis* have been used in other explanatory studies (Arnott and Ruxton, 2002; van Deurs *et al.*, 2009; Lindegren *et al.*, 2018) but are only available with 1-2 years delay, and are therefore of limited value in forecasting recruitment in this stock in an operational setting. We focus our analyses on data that are available with a maximum of a few months delay. An overview of the data employed is provided in Table 1 and the complete time series are found in Figure S1.

**Table 1.** List of all variables considered and the rationale behind. The parameterisation of each variable in the model is also shown, with s() indicates the use of a spline-smoother and other terms indicating the incorporation of that term as a linear response term.

| **Variable** | **Description** | **Rationale** | **Parameterisation** |
| --- | --- | --- | --- |
| **Demographic explanatory variables** |  |  |  |
| *SSB* | Spawning stock biomass | Adult biomass that determines amount of eggs spawned | $s(\log(SSB))$ |
| *N1* | Number of 1-year olds | Number of individuals at age 1 inducing a density dependence | $\log(N1)$ |
| *SumN* | Number of individuals | Entire sandeel population, inducing density dependence | $\log(SumN)$ |
| *TSB* | Total stock biomass | Combination of all of the above | $\log(TSB)$ |
| **Enviromental explanatory variables** |  |  |  |
| *P3* | Jul-Sep temperatures | Temperatures experienced by the adults prior to spawning | *P3* |
| *P4* | Oct-Dec temperatures | Temperatures experienced by the adults prior to / during spawning | *P4* |
| *Q1* | Jan-Mar temperatures | Temperature experienced during egg development | *Q1* |
| *Q2* | Apr-Jun temperature | Temperature experienced by larvae during pelagic drift phase | *Q2* |
| *Q3* | Jul-Sep temperature | Temperature experienced by post-settlement juveniles | *Q3* |
| *Q4* | Oct-Dec temperature | Temperature experienced by post-settlement juveniles | *Q4* |
| **Other explanatory variables** |  |  |  |
| *Cohort* | Cohort year | Included to allow time-variation in the mean productivity of the stock due to systematic shifts in other unquantified variables | $s(Cohort)$ |

### Assessment data

Assessment data used for sandeel modelling is obtained from official ICES advice, based on the stochastic multi-species assessment model, SMS (Pedersen *et al.*, 1999).The SMS model is run in a single-stock mode for sandeel assessments, and integrates data on catches, catch effort, maturity, weight, fishing mortality and natural mortality at a given age (ICES, 2018). All stock assessment data are the current (2021) assessments provided by ICES for area 1r, 2r, 3r and 4 (Figure 2b), where recruits are treated at age 0. From the assessment data, 4 demographic variables are extracted, consisting of spawning stock biomass (SSB), total stock biomass (TSB), number of individuals (SumN) and number of one-year olds (N1). This allows for different types of interactions between the demography, including density dependence and SSB impact on recruitment. All demographic data are log-transformed before use in modelling and converted to log-anomalies (relative to the average log-value over the full time series for each stock).

### Environmental data

High resolution spatial sea surface temperature data is gathered from the Optimum Interpolation Sea Surface Temperature (OISST) product (Banzon *et al.*, 2016). The product is a 0.25° × 0.25° global daily sea surface temperature (SST) data set on a regular grid. Noting that adult sandeel are bound to specific banks (Christensen *et al.*, 2008), we produced daily average temperatures over the banks in each stock area, and then averaged temporally over quarters as follows: P3 and P4 represents the temperature anomalies experienced by the adult sandeel from July to December before and during spawning (i.e. the temperatures experienced by the spawners just before spawning). Q1, Q2, Q3 and Q4 are SST anomalies experienced during the egg, larval and juvenile stages from January to December for a given cohort. All extracted temperatures were converted to anomalies from the average (climatology) over the complete SST time series period (1983 to 2020) prior to use in modelling.

### Models

Here we use generalised additive models (GAM) as the basis for generating predictions, with model variable selection based on a multi-model inference approach. An advantage of the GAM approach is it’s semi-parametric nature that allows for arbitrary but smooth responses. We exploited this feature to incorporated a cohort-based time-varying smoother to allow for changes in the underlying productivity (e.g. due to unquantified variables). This approach allows non-stationarity and systematic shifts in recruitment patterns that would otherwise not be accounted for. For in-depth model descriptions, see supplementary methods.

The total of 11 candidate variables (Table 1) give a total of 2048 possible combinations that could be considered. However, in order to minimize risk of overfitting due to both collinearity between model parameters and the short time-series, predictors are split into three groups (as shown in *Table 1*) based on an exploratory analysis of collinearity (i.e. environmental, demographic and other predictors). Models in the ensemble that incorporated more than one variable in a given group were excluded, giving a total of 819 candidate model structures to be considered.

Following the retrospective-forecasting and time-blocking approach proposed in this framework, (*F*igure *1*), models were first trained on all data up to a cut-off point and the small-sample Aikaike Information Criteria (AICc) calculated and converted to model weights (Anderson, 2008). Each model was then used to predict the distribution of expected recruitment values for each cohort in the second, verification block. The individual model posterior predictions were then combined into an ensemble predictive distribution, with the contribution of each model to the ensemble prediction being determined by the AICc weights. Probabilistic categories (i.e. high, medium and low recruitment) and the expected value (mean across the distribution) were then generated from this ensemble predictive distribution. This process was repeated by moving the cut-off point (forecast issue date) forwards by one year, creating a forecast various lead times (*F*igure *1*b).

We evaluated forecast issue dates from 2007-2020, giving a total of 14 forecasts to evaluate: earlier first-forecast dates struck problems with model stability due to the short time series in area 4 (starting from 1993). We focused on the first forecast (one cohort ahead) here, as this is the most relevant to both the management of the stock and to the associated fishing industry.

### Skill metrics

Multiple performance metrics are used to assess the retrospective forecasts (*Table 2*), including both continuous and categorical skill evaluations (Stow *et al.*, 2009; Jolliffe and Stephenson, 2012; Brun *et al.*, 2016). Continuous skill uses the mean prediction for a root-mean-square error (RMSE) analysis, giving indications of the accuracy of the forecast. Continuous forecasts can use the mean-squared-error skill score (MSESS) to directly compare the forecast with a reference forecast. The categorical forecasts (high, medium and low) are analysed using the hit rate (H), false alarm rate (F) and true skill score (TSS). Using a combination will quantify both the accuracy of the forecast and forecast performance in each tercile (Murphy, 1969).

**Table 2.** Performancel metrics used to evaluate the skill of the forecast system. These values are calculated over all retrospective forecasts at a given lead time. Continuous skill evaluation is also performed for reference forecasts, while the categorical and binary forecast evaluation are only calculated for probabilistic forecasts (Murphy, 1969). MSE contains the mean of the difference between the forecasted (F) and observed (O). The hit rate (H) consists of the proportion of correct forecasts (i.e. true positives (TP) and true negatives (TN)), while false alarm rate (F) is the proportion of incorrect forecasts (i.e. false positives (FP) and false negatives (FN)).

| Name of Forecast Quality Measure | Definition | Range | Application |
| --- | --- | --- | --- |
| Mean square error (MSE) | $\frac{1}{n} \sum_{i=1}^n (F_i - O_i)$ | [0,inf] | Cont. |
| Mean square error skill score (MSESS) | $1 - \frac{MSE}{MSE_{reference}}$ | [-inf,1] | Cont. |
| Root-mean-square error (RMSE) | $\sqrt{MSE}$ | [0,inf] | Cont. |
| Proportion correct / Hit rate | $H = \frac{TP + TN}{(TP + TN) + (FP + FN)}$ | [0,1] | Cat. |
| False alarm rate | $F = \frac{FP + FN}{(TP + TN) + (FP + FN)}$ | [0,1] | Cat. |
| True Skill Score (TSS) | $TSS = H - F$ | [-1,1] | Cat. |

Reference forecasts were selected according to current stock assessment practices: in this way, it was immediately apparent if the forecast outperforms existing procedures. For the sandeel, the official ICES sandeel advice uses either the 10-years moving geometric mean (Area 2r and 4) or the geometric mean of the full time series (Area 1r and 3r) (ICES, 2018). These models are selected as reference forecasts in the MSESS. The skill score ranges from negative infinity to 1, effectively comparing the performance gains from using a given forecast compared to the reference. For categorical forecasts, the reference forecast is selected to be random guessing (33% correct) baseline, both for True Skill Score (TSS) and Ranked Probability Skill Score (RPSS).

### Forecast value

We assess the value of the forecasts using a Richardson cost-loss decision model (Richardson, 2000). Simple economic models, as used here, are widely used in the climate services sector (Pope *et al.*, 2019) to quantify value of e.g. seasonal forecast systems, and provide an intuitive metric for users (Murphy, 1976b). Briefly, the model considers the economic impacts of a particular event that is being forecast (e.g. poor recruitment), and the loss (L) that the user could potentially incur. However, the user also has the ability to avert these losses by implementing precautionary mitigation actions (e.g. based on a forecast), but doing so also incurs a cost (C) (e.g. mothballing processing plants). These two dimensions (i.e. whether the event occurs, and whether the user takes a precaution) each have two outcomes, and therefore form a 2×2 cost matrix (Jolliffe & Stephenson, 2012, see Table 3b). Combining this set of costs with the properties of the forecast system characterised by the contingency matrix (Table 3a) allows the expected expense over the long-term (E) to be calculated when the forecast is always (*E_forecast_*) or never (*E_reference_*) followed. The value (V) of the realised forecast system can then be calculated relative to a perfect forecast system as :

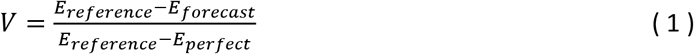

**Table 3.** a) Confusion matrix generated from the retrospective forecasts at a given lead. Constructed from the sum of observed positives (event occurred) and observed negatives (event didn’t occur) with corresponding predicted positives (predict event occurred) and predictive negatives (predicted event not to occur). This results in a matrix of true positive (TP), false positives (FP), false negatives (FN) and true negatives (TN). For recruitment predictions, a TP is when the forecasting system correctly predicts the observed tercile, while a FP is when the system predicts a given tercile, which is not observed. For negative events this is reversed, i.e. FN the tercile is observed while the forecast system doesn’t predict it and TN is when the tercile is not observed and the system doesn’t predict it. b) Cost matrix used to calculate the value of the forecast system. Here a cost (C) is associated with a precaution and a loss is associated with not taking the precaution and the event occurring.

| a) Contingency matrix |  |  | b) Cost matrix |  |  |
| --- | --- | --- | --- | --- | --- |
|  | Observed P | Observed N |  | Event occurs | Event does not occur |
| Predict P | TP | FP | Precaution taken | C | C |
| Predict N | FN | TN | Precaution not taken | L | 0 |

The value of the forecast system, V, is expressed as a non-dimensional number less than 1 and varies as a function of the cost-loss ratio (C/L) of a given user (Richardson, 2000, see eq. 1).

We necessarily extend this analysis to account for the (relatively) small sample size associated with our set of retrospective forecasts and therefore estimate the uncertainties in the value. We model the retrospective contingency table (Table 3a) using a Bayesian multinomial model implemented in Stan (Stan Development Team, 2020) to estimate the vector of true probabilities ***p*** = {*p_TP_*, *p_FP_*, *p_FN_*, *p_TN_*} of each quadrant of the contingency table. The posterior predictive distribution of ***p*** was then sampled 4000 times and used to construct a corresponding large set of contingency tables and therefore the statistical distribution of the forecast system value, V.

## Results

Assessment of the predictions is presented at a forecast lead of one cohort beyond the final year of the assessment, mimicking potential operational usage in these stocks. We find that the stocks in area 1r and 2r have the highest continuous forecast accuracy, while areas 3r and 4 show higher RMSEs (Figure 3a): this dichotomy closely parallels the lengths of the time series of each area (areas 3r and 4 being appreciably shorter) and we hypothesis that the reduced amount of training data may limit the forecast skill. Furthermore, the assessment of area 1r is widely perceived as being the most reliable of the four: the poor performance in areas 3 and 4 in particular may be due to the poor quality of the assessment as much as the poor quality of the forecast. The portfolio forecast, on the other hand, has the highest overall accuracy, showing that the aggregation of predictions can lower the RMSE, highlighting the smoothing effect associated with aggregating noisy data sets.

**Figure 3.**
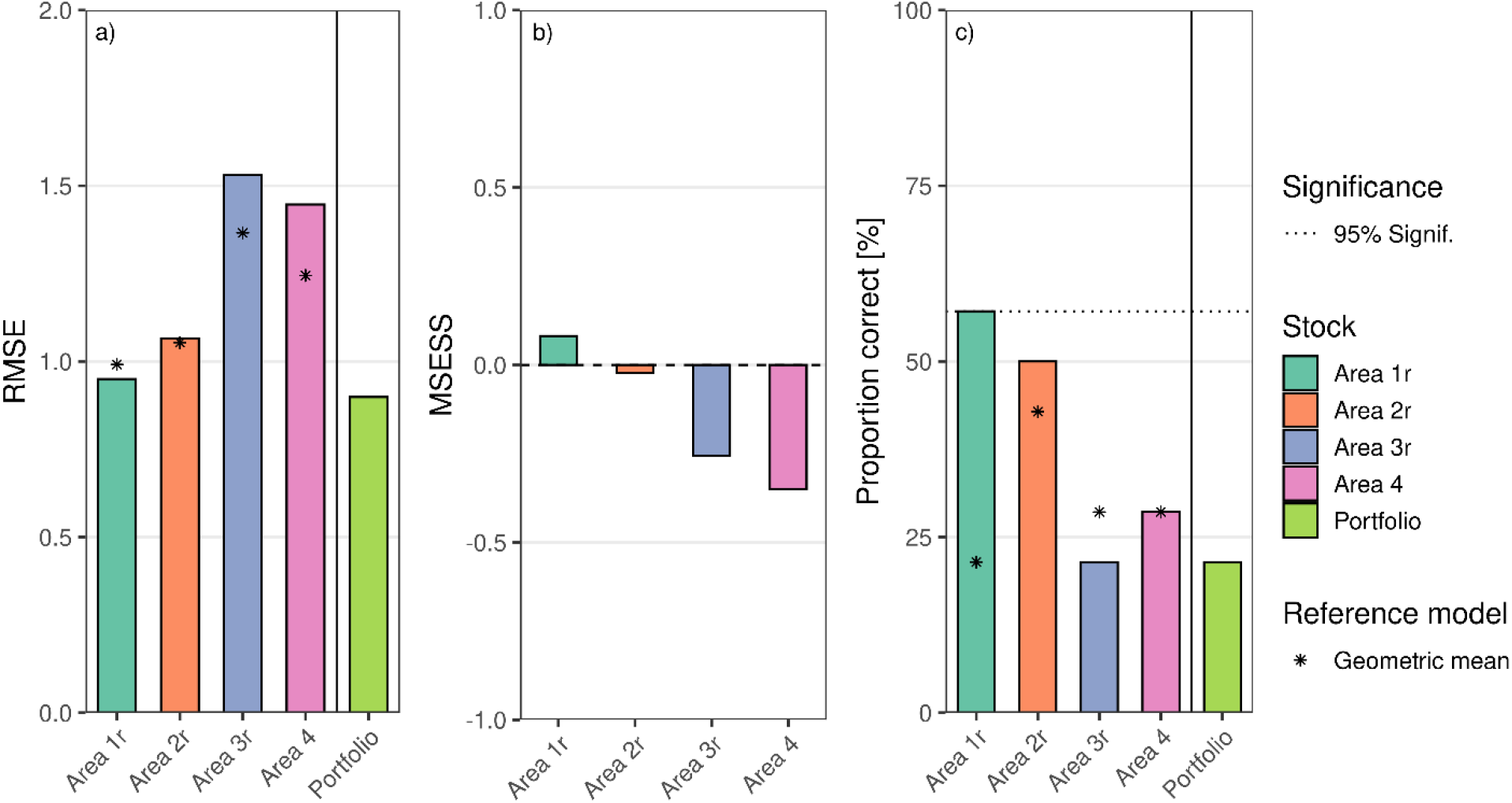
Recruitment forecasts outperform reference forecasts in some cases. a) Root-mean-squared error of the different management areas and the portfolio forecast at lead 1. Area 1r and 2r show highest accuracy of individual forecasts, while the portfolio is the overall most accurate, indicating the presence of the portfolio effect. Stars show the reference geometric mean RMSE for the individual management areas. b) Mean-squared error skill score of the individual forecast products for lead 1. Official recruitment prediction model is used as a reference model. Here area 1r and 2r shows better or equal performance to the reference models, while area 3 and 4 has a negative skill score. c) Hit rate of the different management areas indicating the percentage of correct retrospective forecasts at lead 1. Dashed line indicates the 95^th^ percentile level of the random guessing reference forecast. Area 1r are significantly better than random guessing at 57% hit rate, while area 2r are borderline significant with 50% hitrate. Area 3r, 4 and the portfolio shows large drop-offs in hit rate with hit rates below 30%. Stars show the reference geometric mean hit rate.

Comparing our forecasts against the existing models used in the assessment of this stock (geometric mean) places their skill in context. In management area 1r, the continuous forecast accuracy is better than these reference models (Figure 3a), giving a positive mean-squared error skill score (MSESS) (Figure 3b). The performance of Area 2r is on a par with the reference model, while area 3r and 4 both show a negative MSESS at lead 1, indicating that the forecast model ensemble would not be an improvement over the geometric mean reference model when used as a continuous forecast.

The categorical performance of the forecast models is also broadly similar. Hit rate metrics (how often the system correctly forecasts high, medium or low recruitment) also shows best results in management area 1r, with 57% correct (Figure 3c), outperforming the outperforming expectd 33% correct associated with the random guessing of terciles (p=0.02, one-tailed test). Area 2r sees a hit rate of 50% correct (p=0.06, one-tailed test), significant at the 90% level. A large drop off in hit rate is seen in area 3r and 4 (respectively at 21% and 28%), where performance is not significantly better than random guessing (p=0.74, and p=0.52, one-tailed tests). The portfolio categorical forecast, on the other hand, performs poorly and is not significantly better than random guessing at a 21% proportion correct (p=0.74, one-tailed test): while aggregating improves the performance of continunous forecasts, it clearly deteriorates categorical forecasts.

Further insight into the forecast system can be gained by examining the skill of predicting individual terciles. The true-skill score (TSS) metric combines the specificity (true-positive rate) with the sensitivity (true-negative rate) for a categorical forecast and is applied here to each tercile in turn. The TSS indicates area 1r being the only management area where the model can reliably differentiate all three categories (Figure 4), consistently outperforming random guessing (i.e. where TSS=0). Most other areas do not show a significant ability to differentiate, either due to the small sample size or poor model skill. For example, area 2r’s TSS is not significantly different from zero for all terciles, in part due to the relatively low recruitment seen in the stock in recent years, affecting the ability of the TSS metric to quantify the forecast skill. Areas 3r and 4 have negative or zero skill scores in all categories, likely due to the aforementioned poor quality of these assessments propagating into these forecasts and resulting in a wide prediction distribution. The portfolio forecast shows similar TSS values to area 2r, with no categories reaching levels where the system is able to correctly distinguishing between terciles.

**Figure 4.**
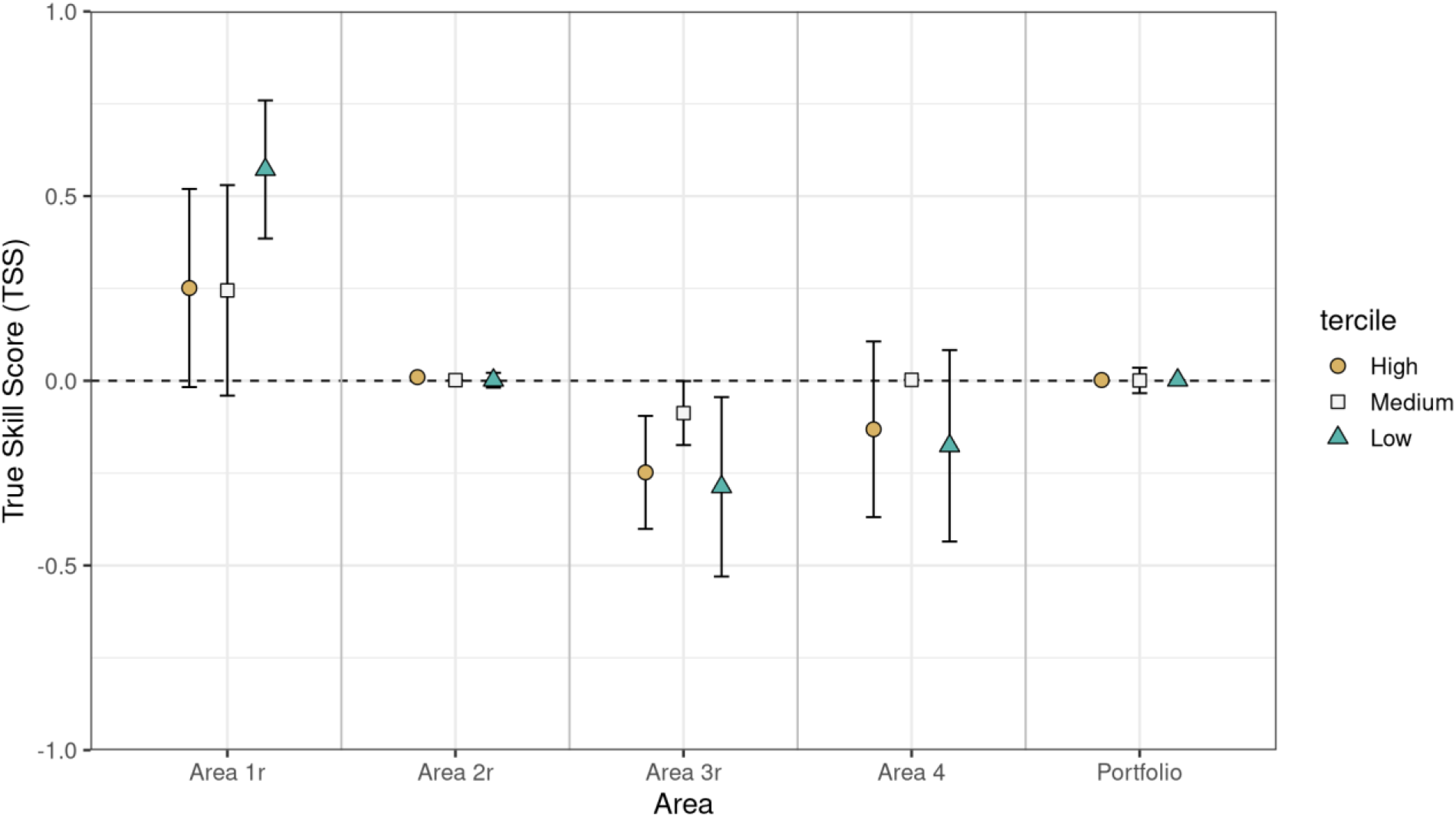
Categorical recruitment forecasts show skill in some areas. Model skill at lead 1 is represented as the True (Peirce) Skill Score (TSS), which ranges between +1 and −1, and has a value of 1 for perfect skill, and 0 for random guessing (black dashed line). Negative values indicates perverse forecast. The 95% confidence interval for the estimated skill score are shown as error bars on each of the points. Recruitment stocks are shown, with shapes indicating the corresponding recruitment tercile. A positive TSS is seen for all recruitment terciles in area 1r, while all other models shows utility close to or worse than random guessing.

We assessed the value of the forecast for all areas. The cost-loss decision model for Area 1r (Figure 5) shows positive values in all forecast categories, with especially the low recruitment prediction showing the highest value over a broad range of cost/loss ratios (Figure 5a). All categories peak at a cost-loss ratio of 0.33, as is expected from theoretical analyses of this model (Jolliffe and Stephenson, 2012). We account for the small sample size and propagate the uncertainty that it creates into the forecast value by estimating the probability of a positive expected value for a given cost/loss ratio (Figure 5b): this metric provides decision makers with an indicator when using the forecast will lead to a positive economic return. Here the peak is still seen at a cost-loss ratio of 0.33, where all categories have above 65% probability of a positive expected long-term value. Following the low recruitment forecast for this cost-loss ratio (i.e. 0.33) will result in a 96% probability of positive value from the forecast, but probabilities above 50% are also seen across a wide range of cost-loss ratios. While area 2r, 3r and 4 generally can’t provide the same levels of value, area 2r could prove valuable when following the high forecast (Figure S3).

**Figure 5.**
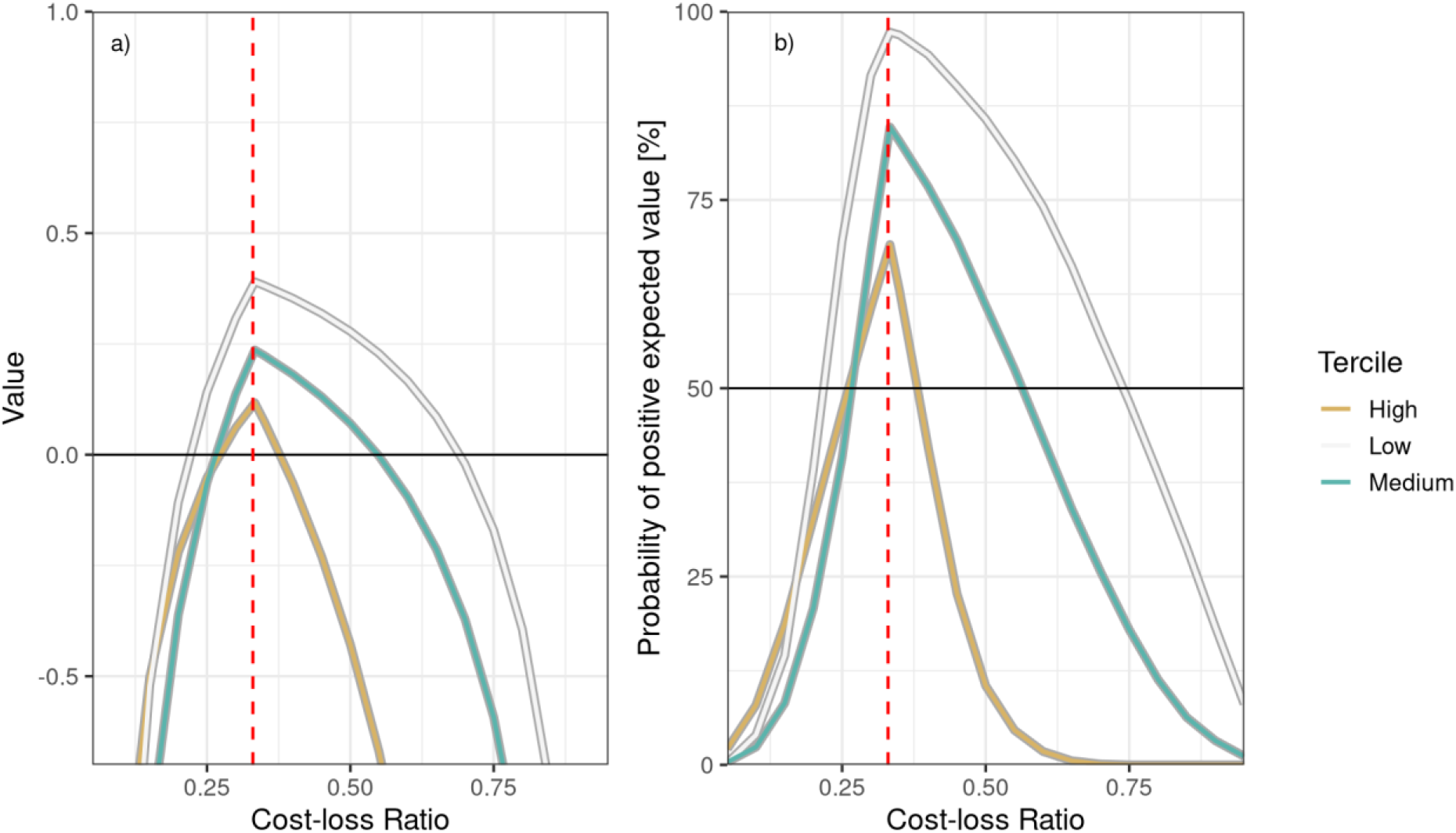
Positive economic value is seen in area 1r recruitment forecasts. Long-term value of a cost-loss decision model in area 1r, simulated from a multinomial confusion matrix model. a) Tercile divided value given cost-loss ratios. Solid line indicates zero value. Positive value is seen in all terciles, peaking at a cost-loss ratio of 0.33. Most value can be gained by following the low tercile forecast, which corresponds with the highest observed TSS. b) Tercile divided probability of a positive expected value. Calculated from a Bayesian posterior distribution, indicating the probability of drawing a positive value at a given cost-loss ratio. Peak probability is seen at cost-loss ratio of 0.33, where all terciles shows above 65% probability of a positive expected value.

## Discussion

Here we present a framework for robustly assessing the skill and value of recruitment predictions in a way that is relevant to their use in an operational setting. The case study that we have examined, for four sandeel stocks in the North Sea, illustrates several important conceptual points that deserve particular attention.

Firstly, we show the importance of assessing a forecast system with multiple metrics. While in-sample performance and explanatory metrics are good for finding correlations (and thereby highlighting possible causality), the assessment of predictive skill is quite different and should primarily be shaped by the needs of the forecast user. For example, we identify an overall high forecast accuracy in area 2 (RMSE in Figure 3), but the ability to distinguish between the two lower terciles is poor (Figure 4). Stock assessors may focus on the MSESS as a criteria for uptake, while industry might be more interested in performance in a specific category (e.g. ability to forecast poor year classes) or long-term economic value. Furthermore, the value of a forecast to users within the same sector (e.g. two different fish processing plants) may differ due to differences in their underlying risk profile (i.e. cost-loss ratio) such that while the forecast system may be advantageous for one user, it may not be of use to another. Understanding the decision-making needs of the user is therefore essential to the production of a good forecast (Murphy, 1993; Payne *et al.*, 2017).

While the application of the cost-loss model to estimate forecast value has a clear interpretation in a commercial context, it is less clear how relevant this approach is to fisheries management. Here, cost-loss decision models encapsulate both the costs and losses associated with correctly and incorrectly forecasting recruitment. These ideas can be relevant to fisheries management, as the managers can use this knowledge as the basis of the forecast evaluation, assigning value on e.g. true positives versus false positives. This allows managers and users to understand how the forecast can be incorporated, and how forecasts can and should be used in the management of a given stock. While not an economic gain, the value metric of a forecast can be used to assess and manage stocks sustainably, providing the managers with the tools to properly assess how to incorporate forecasts into decision making.Our demonstration of the framework here is based on the use of recruitment estimates directly from the stock assessment, as is still common in the field. It is nevertheless important to remember that these data are estimates that are also uncertain (Brooks and Deroba, 2015). The framework presented here has the ability, however, incorporate a more robust treatment of such uncertainties. For example, uncertainty estimates (e.g. in recruitment) can be incorporated directly into forecast model if desired. Retrospective biases in the stock assessment incorporated into the model fitting procedure by e.g. fitting the forecast model to stock-assessment outputs based on a model up to 2007, and then predicting forward in time from there. While such an approach would be idealogically cleaner, it was not possible here due to technical challenges in producing a sufficient number of retrospective assessments for these stocks. A further extension would be to incorporate the recruitment forecast model directly into the stock assessment model, thereby making a seamless assessment and recruitment prediction system. Regardless of the approach, the framework presented can adapt to both the technical limitations of the system being studied, and changing norms in the approach to this issue.

Finally, our results for North Sea sandeel show that our understanding of recruitment predictability needs to be re-assessed. Contrary to the wide-spread belief that recruitment can’t be forecast, we have shown in a setting that directly mirrors operational useage that skilful and valuable recruitment forecasts can be made. Shifting the way that we assessment recruitment skill from an explanatory to predictive setting greatly increases the confidence in, and transparency of, these results, and paves the way for their direct up-take in decision making. Furthermore, taking the next step of assessing the value of these forecasts gives a more nuanced view that is directly relevant to decision-makers, particularly in the commercial sector. These results therefore open the way for a new paradigm in addressing this long-running, but fundamental question in fisheries management.

## Acknowledgements

The research leading to these results has received funding from the European Union’s Horizon 2020 research and innovation programme under grant agreement No 727852 (Blue-Action).

This study was co-funded the PANDORA project which received funding from the European Union’s Horizon 2020 research and innovation programme under the grant agreement no. 773713.

## Data availability

The data that support the findings of this study are available from the corresponding author upon reasonable request.

