## Supplementary material for "A framework for assessing the skill and value of operational recruitment forecasts"

Running header: Skill & value in recruitment forecasting

### **Supplementary materials**

#### **Model description**

The multi-model inference (Burnham and Anderson, 2004) module is developed using R and combines the usage of generalized additive models (GAMs) (Hastie and Tibshirani, 1986) with the multi-model inference philosophy and ensemble modelling. A global sandeel GAM model is selected from ecological knowledge and data availability (see eq. 1 and Table 1). From this global GAM a subset of models is created and weighted according to AICc (The small sample Akaike Information Criteria, which allows for smaller sample sizes in relation to model parameters (Anderson, 2008)).

$logRecruits \sim s\left( logSSB \right)+s\left( Year \right)+logSumN+logN1+logTSB\ldots$( 1$)$

$$P3+P4+Q1+Q2+Q3+Q4$$

All demographic and environmental variables are included in this global model, including a cohort year with a non-parametric smoother, allowing for time variance in the baseline productivity to be incorporated (e.g. due to unquantified variables). To minimize risk of overfitting due to collinearity between model parameters, predictors are split into groups based on correlations (e.g. temperatures and demographic predictors). For the sandeel a total of 819 candidate models are selected from the global model, creating an ensemble using all subset models to predict. Each model in the subset will be limited to only include one parameter from each group (see eq. 2 for an example of an allowed subset model).

$logRecruits \sim s\left( \mathrm{logSSB} \right)+\text{s(Year) + logN1 + Q1}$	( 2 )

The subset of models is weighted according to AICc values using the relative likelihood of each model (see eq. 3), where the weight of a model is determined by the models relative likelihood divided by the sum of relative likelihoods in the ensemble (Anderson, 2008). This leads to a model ensemble, where models with lowest scoring AICc are given highest weight, resulting in the largest contributions to the prediction, using the ranking of the models to determine influence. The large subset of weighted models will be used to make ensemble predictions, creating a probabilistic predictive distribution from the fitted models. The predicted probabilities of the forecasted cohort year class strength are compared to three historical terciles (high, medium or low recruitment), where a categorical forecast is chosen from the most likely tercile. Tercile thresholds are based on the values from the complete time series.

$w_{i}=\frac{e^{-0.5*{AIC}_{i}}}{\sum_{j=1}^{N} e^{-0.5*{AIC}_{j}}}$	( 3 )

#### **Model Results**

Our results parallel those seen in other sandeel recruitment studies (Arnott and Ruxton, 2002; van Deurs *et al.*, 2009), where relative variable importance shows SSB is not a primary recruitment driver (Figure S2). Furthermore, analysis of the retrospective models shows a clear shift in recruitment dynamics over the evaluated period, indicating that the system is non-stationary (Figure S2). For the lesser sandeel additional drivers has been shown to influence recruitment, such as prey concentrations (van Deurs *et al.*, 2009) and circulation patterns (Henriksen *et al.*, 2018). The current framework is able to include such types of drivers, allowing for specialised forecasts for a given species or stock and basing it on already known drivers. However, operational usage does require data availability at the time of forecasting, limiting the options in selection of potential predictors.

### **Supplementary figures**

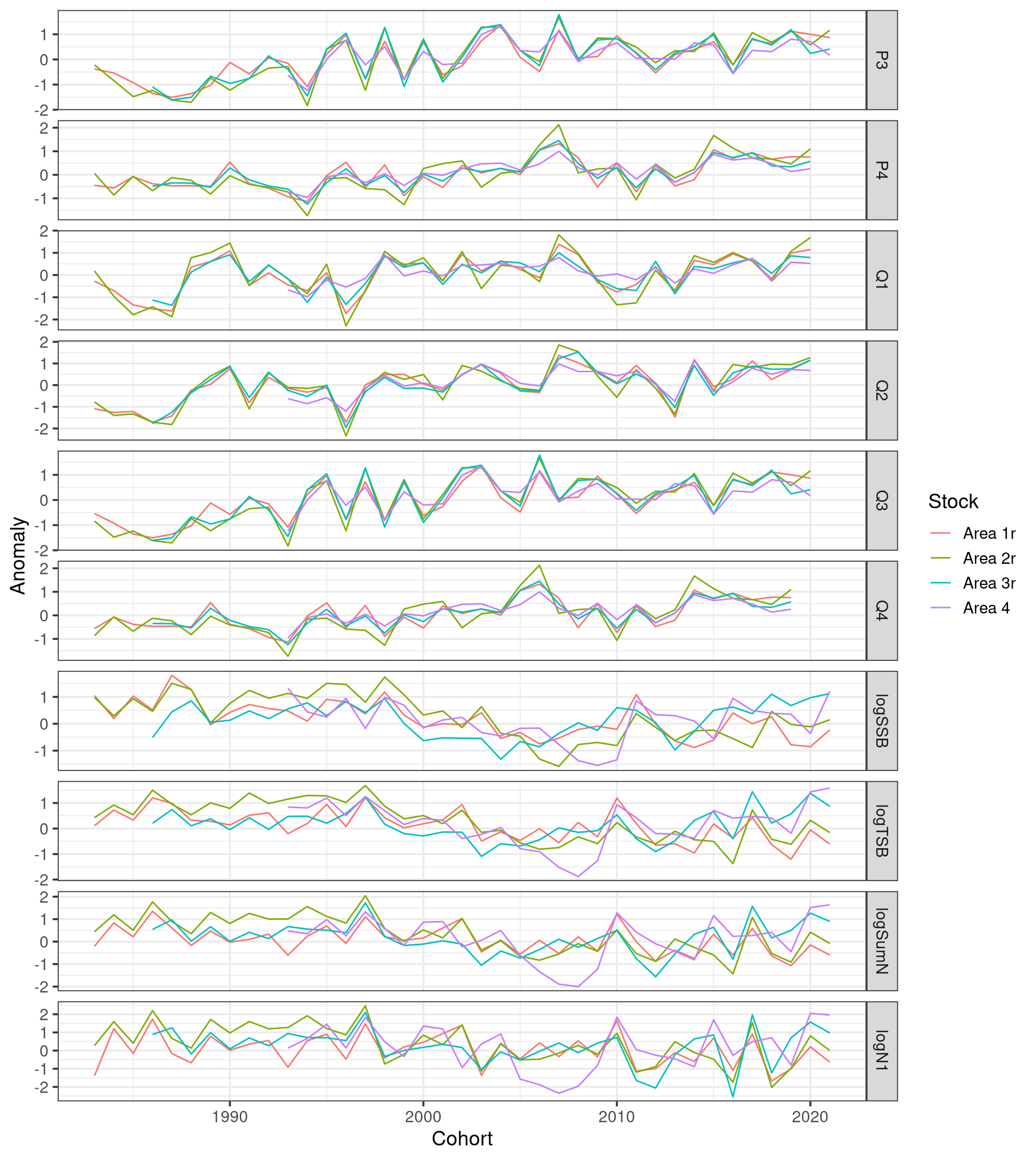

Supplementary figure 1 All data used for modelling. Contains demographic information from spawning stock biomass (logSSB), total stock biomass (logTSB), number of individuals (logSumN) and number of 1-year olds (logN1). Environmental information consists of sea surface temperature (SST) from the two quarters before spawning and the 4 quarters after. All modelling data are treated as anomalies.

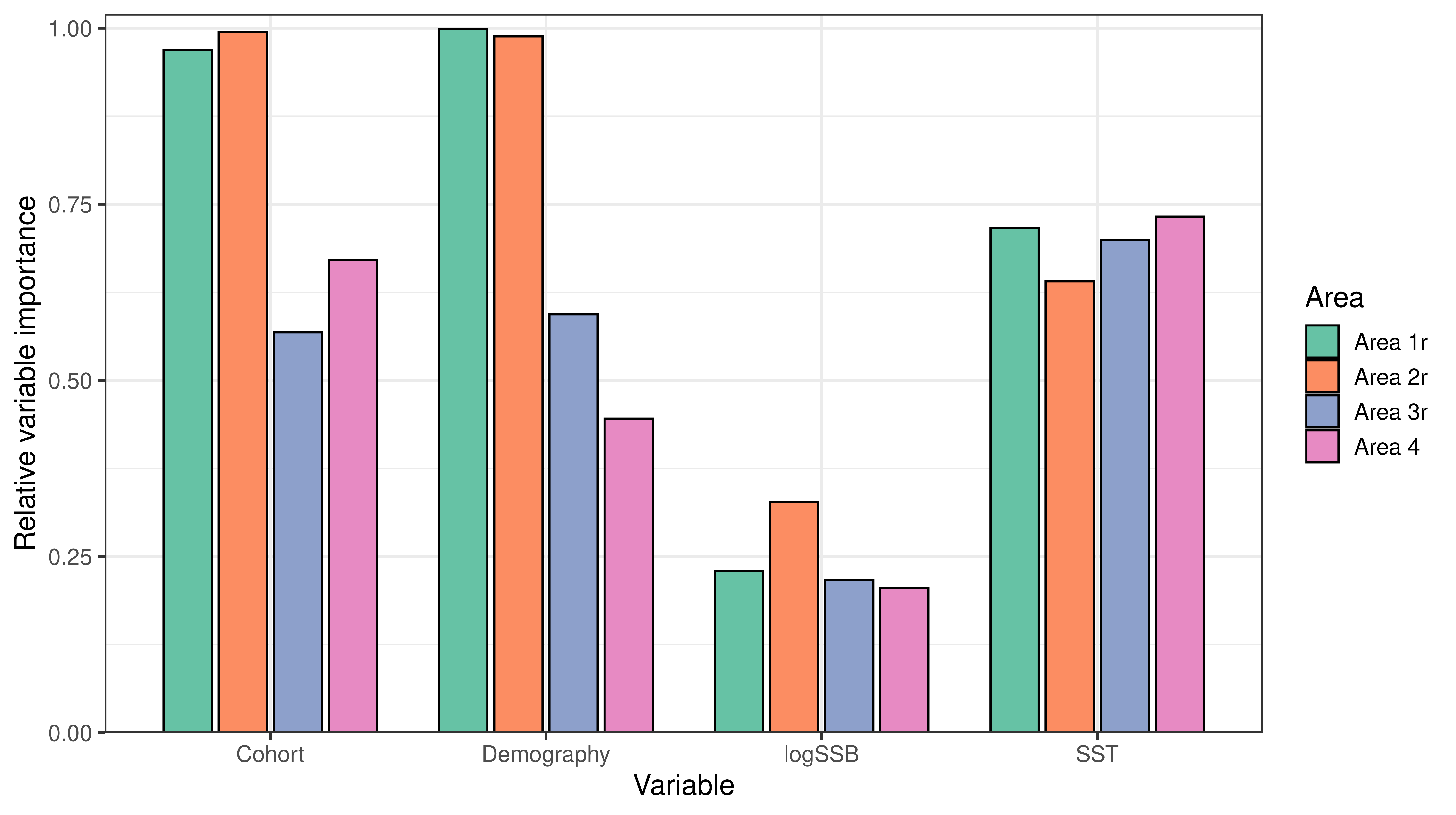

Supplementary figure 2 Relative variable importance of variables in the multi-model ensemble. Here the SST and non-SSB demographic variables (as classified in Table 1) are grouped together, with the exception of the spawning stock biomass. The main predictive variable in most areas is the non-parametric Cohort parameter. Here, SSB are seen having a low impact on the overall modelling ensembles across the areas, where other demographic variables and the SST is contributing more.

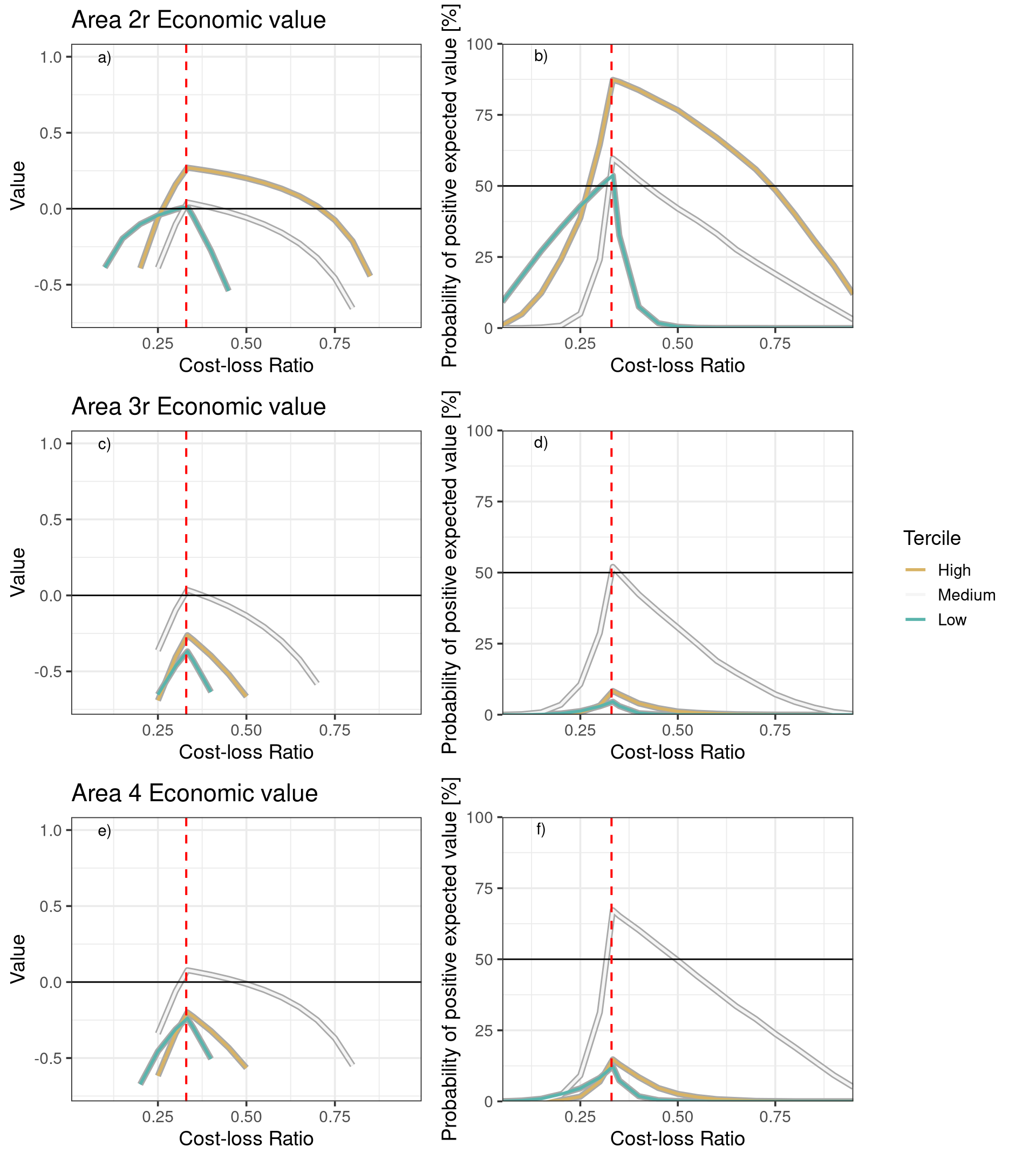

Supplementary figure 3 Long-term value of a cost-loss decision model in areas 2r, 3r and 4, simulated from a multinomial confusion matrix model. a, c, e) Expected forecast value as a function cost-loss ratios for each tercile. Solid line indicates net- zero value. b, d, f) Probability of a positive expected value by tercile. Solid horizontal line indicates equal probabilities of a net gain or loss from using the forecast. Calculated from a Bayesian posterior distribution, indicating the probability of drawing a positive value at a given cost-loss ratio. Area 2r shows positive expected value in the high tercile, while the other areas show very low probability of expected positive value.
